# Improving laser standards for three-photon microscopy

**DOI:** 10.1101/2020.09.09.289603

**Authors:** Deano M. Farinella, Arani Roy, Chao J. Liu, Prakash Kara

## Abstract

**Significance:** Three-photon excitation microscopy has double-to-triple the penetration depth in biological tissue over two-photon imaging and thus has the potential to revolutionize the visualization of biological processes *in vivo*. However, unlike the ‘plug-and-play’ operation and performance of lasers used in two-photon imaging, three-photon microscopy presents new technological challenges that require a closer look at the fidelity of laser pulses.

**Aim:** We implemented state-of-the-art pulse measurements and developed new techniques for examining the performance of lasers used in three-photon microscopy. We then demonstrated how these techniques can be used to provide precise measurements of pulse shape, pulse energy and pulse-to-pulse intensity variability, all of which ultimately impact imaging.

**Approach:** We built inexpensive tools, e.g., a second harmonic generation frequency resolved optical gating (SHG-FROG) device, and a deep-memory diode imaging (DMDI) apparatus, to examine laser pulse fidelity.

**Results:** First, SHG-FROG revealed very large third order dispersion (TOD). This extent of phase distortion prevents the efficient temporal compression of laser pulses to their theoretical limit. Furthermore, TOD cannot be quantified when using a conventional method of obtaining the laser pulse duration, e.g., when using an autocorrelator. Finally, DMDI showed the effectiveness of detecting pulse-to-pulse intensity fluctuations on timescales relevant to three-photon imaging, which were otherwise not captured using conventional instruments and statistics.

**Conclusions:** The distortion of individual laser pulses caused by TOD poses significant challenges to three-photon imaging by preventing effective compression of laser pulses and decreasing the efficiency of nonlinear excitation. Moreover, an acceptably low pulse-to-pulse amplitude variability should not be assumed. Particularly for low repetition rate laser sources used in three-photon microscopy, pulse-to-pulse variability also degrades image quality. If three-photon imaging is to become mainstream, our diagnostics may be used by laser manufacturers to improve system design and by end-users to validate the performance of their current and future imaging systems.

## 1 Introduction

The now ubiquitous field of two-photon microscopy has been used to visualize a wide array of biological phenomena *in vivo*, from imaging neurons in healthy animals^1-3^ and in disease models such as Alzheimer’s^4^, to imaging the movement of fluorescently-labeled viruses invading the lungs^5, 6^ and the mechanisms of glomerular filtration in the kidneys^7^. Three-photon microscopy builds on the success of two-photon microscopy by using longer excitation wavelengths to reduce light scattering and a three-photon absorption process to achieve better nonlinear confinement in biological tissue, resulting in higher signal to background ratio at larger depths^8-11^. However, three-photon excitation also poses unique challenges due to its heightened sensitivity to the fidelity of the laser source.

Since three-photon excitation requires the near-simultaneous absorption of three photons, laser pulses with higher photon density (higher energy and shorter pulse duration) are used when compared to two-photon microscopy. As a result of using higher energy pulses, lower repetition rates must be used to manage the amount of heat delivered to the sample. Consequently, the integrated fluorescence signal used to construct imaging pixels is a result of fewer excitation pulses. Thus, pulse-to-pulse intensity fluctuations on the timescale of microseconds to milliseconds to seconds will degrade the imaging quality of biological samples. Additionally, because the three-photon absorption rate is proportional to intensity cubed^12^, distortion of individual pulses in the excitation source can detract from excitation efficiency. Therefore, examining time-dependent biological processes that produce meaningful changes in fluorescence brightness requires a stable, distortion-free three-photon excitation source.

## 2 Methods

### 2.1 Simulating measurements of laser pulses with various phase distortions

Gaussian shaped laser pulses were analytically defined with no phase distortions, group delay dispersion (GDD), third order dispersion (TOD), and a combination of the two to show the resulting shape of their electric field and temporal intensity envelope shown in Fig. 1(a)-1(d), and Fig. 1(e)-1(h) respectively. To explore the differences in measuring these laser pulses with different techniques, autocorrelation and second-harmonic generation frequency resolved optical gating (SHG-FROG) measurements were then simulated by calculating the analytical intensity autocorrelation, and computing the SHG spectrogram in Fig. 1(i)-1(l), and Fig. 1(m)-1(p) respectively.

**Fig. 1.**
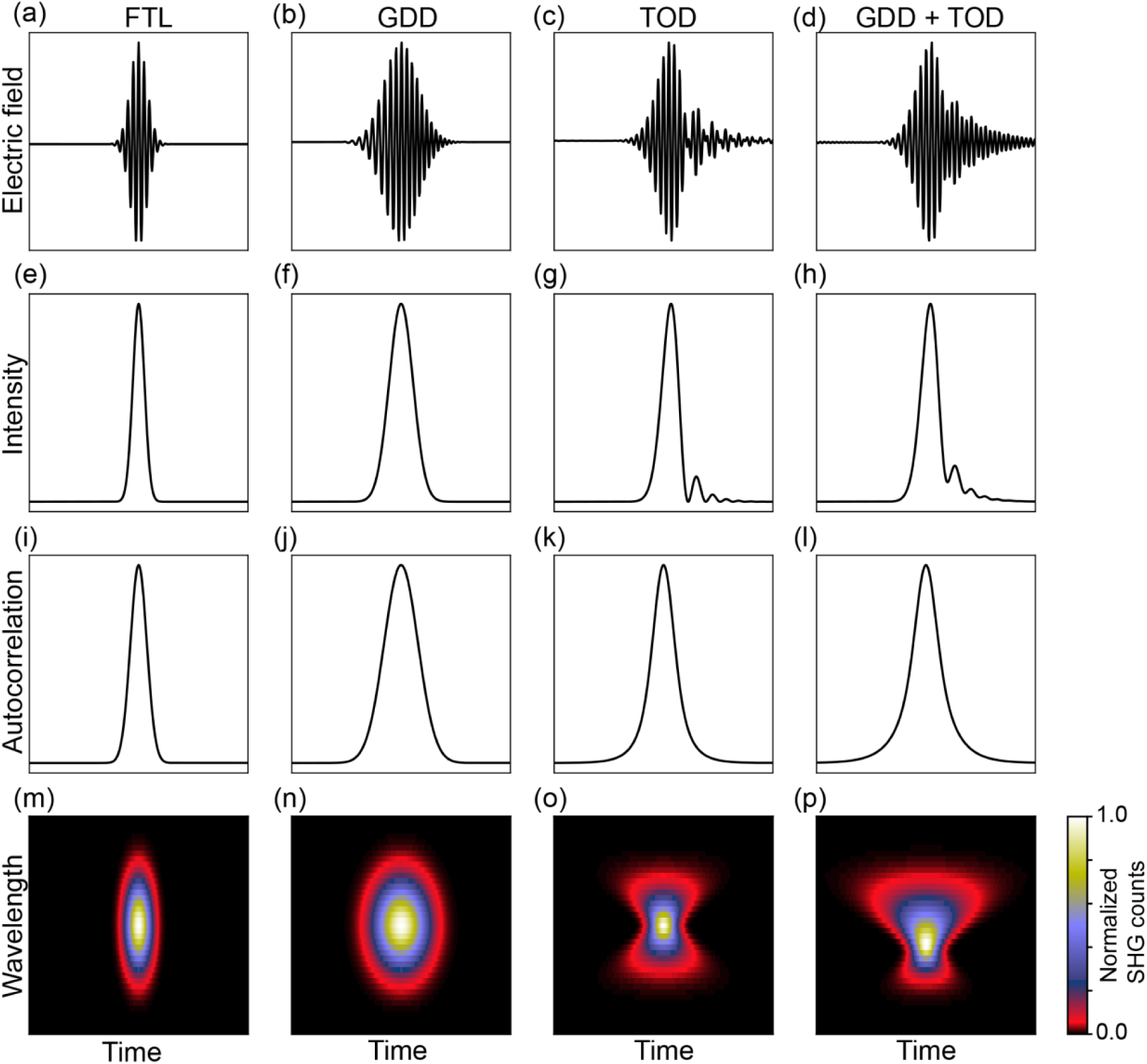
Simulations of distortions of an ideal Gaussian pulse shape caused by GDD and/or TOD. (a-d) Electric field. (e-h) Laser temporal intensity envelope. (i-l) Simulated intensity autocorrelation. (m-p) SHG spectrogram. (a,e,i,m) Laser pulses at the FTL display an ideal pulse shape, where the instantaneous wavelength of the electric field is fixed and the temporal intensity envelope is minimized, leading to a Gaussian shaped intensity autocorrelation and elliptical SHG spectrogram. (b,j,f,n) Laser pulses with GDD show a change in instantaneous wavelength of the electric field as a function of time, and a broadening of the temporal intensity envelope, leading to a broadened Gaussian intensity autocorrelation and elliptical SHG spectrogram. (c,g,k,o) Laser pulses with TOD show a beating of the electric field of the laser pulse and a modulated temporal intensity envelope outside of the central peak, resulting in an intensity autocorrelation of a different shape and an SHG spectrogram that takes on an hourglass shape. (d,h,l,p) Laser pulses with both GDD and TOD exhibit a combination of these effects leading to an intensity autocorrelation with accentuated wings and an hourglass shaped SHG spectrogram with an asymmetric broadening due to the GDD.

### 2.2 Measuring the duration of laser pulses for three-photon microscopy

Laser pulses from a 1 MHz Spirit-NOPA (set at 1300 nm central wavelength, Spectral Physics) were characterized by a home-built scanning SHG-FROG device^13^ [see Supplementary Fig. S1]. The laser pulses were attenuated by uncoated fused silica wedges (PS811, Thorlabs) and further attenuated by an achromatic half-wave plate (AHWP05M-1600, Thorlabs) and SF2 polarizing beam splitter (05FC16PB.9, Newport). The leftover white light in the NOPA laser profile was cleaned with a long-pass filter (FELH1150, Thorlabs) to remove beam profile pollution and simplify alignment. For FROG measurements of the entire width of the laser beam, the beam was down-collimated by a 2x C-coated telescope (GBE02-C, Thorlabs). After down collimation and attenuation, the laser beam was aligned into the entrance irises of the SHG-FROG with gold mirrors (10D20ER.4, Newport).

In the SHG-FROG device, the laser pulses were split into two beam paths (a reference path, and a delay path) by a pellicle beam splitter (PBS-2C, Newport). The delay path had a variable path length by varying the position of a retro-reflecting mirror pair on a delay stage. The laser pulses from each path were steered by gold mirrors (10D20ER.4, Newport) onto the surface of a f = 50 cm mirror (20DC1000ER.2, Newport), which spatially overlapped the laser pulses in a type I β-Barium Borate crystal (BBO, United Crystal). The temporal overlap was achieved by varying the length of the delay path to generate SHG. The SHG signal was then directed by a mirror (10D20ER.2, Newport) through a B-coated f = 10 cm lens (LA1509-B, Thorlabs) to eventually focus into the slit of a spectrometer (Flame-S-VIS-NIR, Ocean Optics).

FROG traces were taken by spectrally resolving the SHG signal from laser pulses overlapped in time and space inside the BBO crystal. Custom LabVIEW software was used to send commands to the motorized actuator (CONEX-LTA-HS, Newport) to perform the scan and record the spectrum at each delay point, which constructed the FROG trace/spectrogram. We took 201 spectra spaced by ∼ 1.9 fs to create the spectrogram.

### 2.3 FROG phase-retrieval algorithm

Data taken with the scanning SHG-FROG was then processed to retrieve the spectral phase^14^. The retrieval of the spectral phase from the measured SHG spectrogram allows for the determination of the underlying shape of the laser pulse temporal intensity envelope [e.g., Fig 2(a)-2(c)]. Data was extracted from the measured spectrogram in the 600–700 nm wavelength range, centered at the second harmonic, i.e., at half the central wavelength of the laser. Then the data was padded with zeros at 50% of the full width and ultimately binned to a grid of 256 × 256 pixels. The phase retrieval algorithm was executed on the binned data until a FROG error of g = 6.9 ×10^−3^ was reached.

**Fig. 2.**
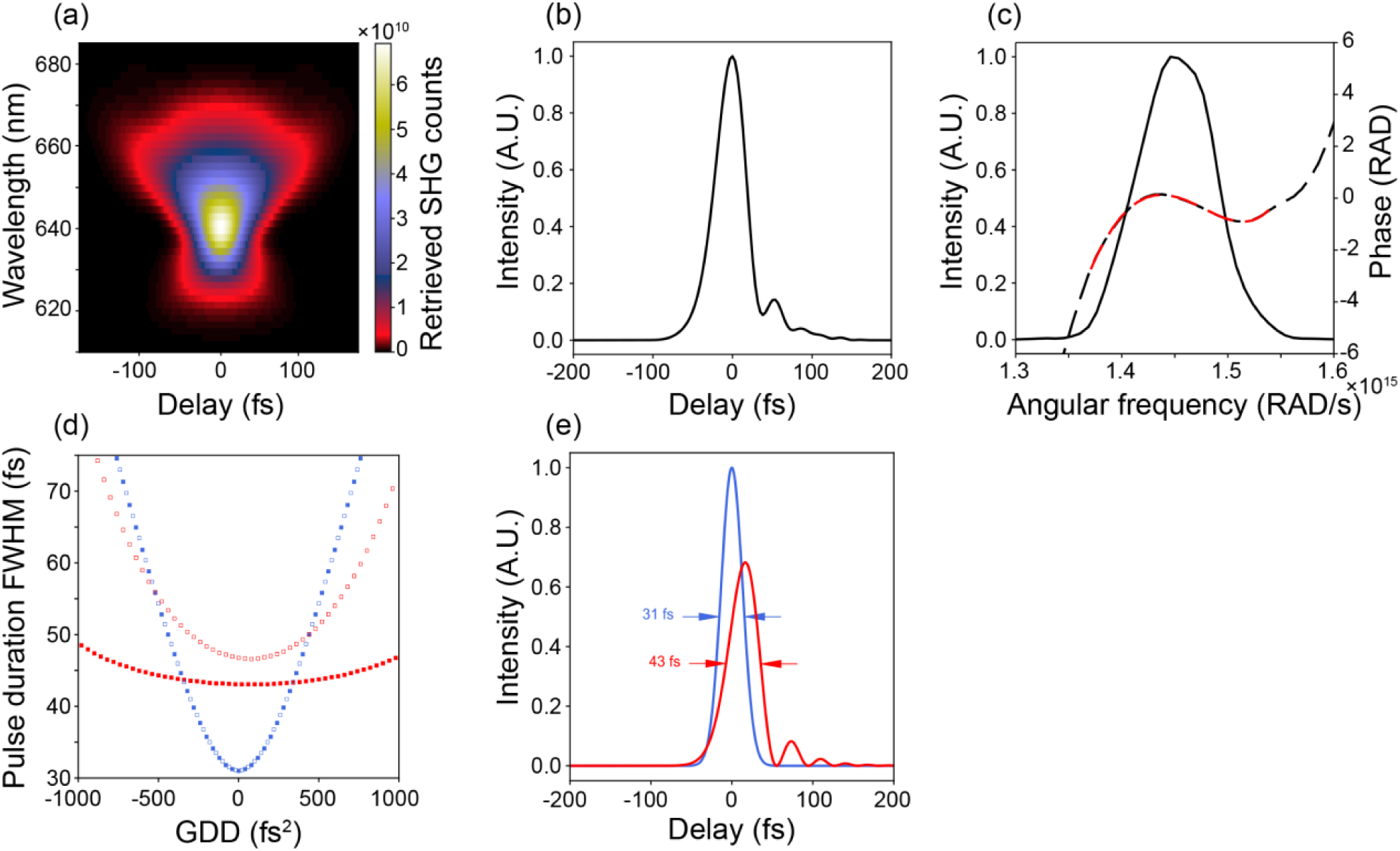
Measurements of spectral intensity and phase by SHG-FROG and modeling of laser pulses. (a) The retrieved SHG spectrogram. (b) The temporal intensity envelope. (c) Spectral intensity and phase. The black solid line represents the laser spectral intensity, the dashed black line represents the spectral phase, and the dashed red line represents a polynomial fit to the measured phase. (d) Analytical modeling of the effects of TOD on pulse compression and measurement by autocorrelation. Data shown as red squares shows the effects of adding or removing GDD from Gaussian laser pulses with the TOD measured from our laser source (∼ 22,750 fs^3^). Data shown as blue squares shows the effects of adding or removing GDD from Gaussian laser pulses with no TOD. Filled squares represent the actual pulse duration and unfilled squares are the pulse duration from a simulated autocorrelation measurement. (e) Modeling the effects of TOD on the temporal intensity envelope. A comparison of the temporal intensity envelope of a Gaussian laser pulse with the TOD measured from our laser shown in red and a Gaussian laser pulse with no TOD shown in blue, assuming GDD = 0 fs^2^.

### 2.4 Determination of spectral phase coefficients from FROG measurements

The measured spectral phase as a function of frequency [e.g. Fig. 2(c)], can be approximated by a Taylor series expansion with respect to the central frequency of the laser spectrum *ω*_0_ to obtain the following Eq. 1.

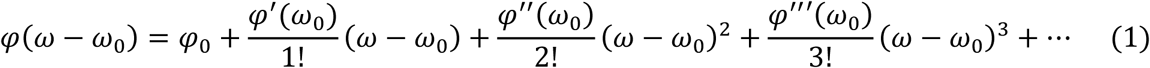

Here *ω* is the angular frequency, and *φ* is the spectral phase. The zeroth order term and first order term result in a shift in the carrier envelop phase and a group delay respectively, and consequently do not contribute to changes in pulse width. The quadratic term is known as the group delay dispersion (GDD) and is the first term in the expansion that results in pulse broadening. GDD can be compensated with many common pulse compressor designs (i.e., prisms, diffraction gratings, bulk material), and is defined as, GDD ≡ *φ*′′(*ω*_0_). Phase coefficients beyond GDD are typically referred to by their order [i.e., third order dispersion (TOD) ≡ *φ*^′′′^(*ω*_0_), and fourth order dispersion (FOD) ≡ *φ*^′′′′^(*ω*_0_) etc.]. These higher order terms are generally costly and difficult to compensate in the required amount while simultaneously compensating for GDD, especially for the high repetition lasers used in multi-photon imaging.

To determine the GDD and TOD from the FROG measurement, the measured spectral phase was fit by a fourth order polynomial centered at the peak wavelength of the retrieved spectrum [e.g., Fig. 2(c)]. The fit extended to where the retrieved spectral intensity fell to ∼ 5% of the peak intensity and was used to compute the GDD and TOD coefficients as described in Eq. 1. Due to the arrow of time ambiguity in SHG-FROG, the sign of the GDD and TOD was determined empirically by introducing increments of a material with a known GDD and TOD into the path and taking additional SHG-FROG measurements to determine if the GDD and TOD increased or decreased with the introduction of the material. This approach revealed the sign of GDD to be negative, and the sign of TOD to be positive.

### 2.5 Analytical modeling of the effects of GDD and TOD on laser pulses

To analytically model the effects of GDD and TOD on the temporal intensity envelope of our laser pulses, we first measured the spectrum of the Spirit-NOPA laser beam using a spectrometer (NIRQUEST, Ocean Optics). This spectrum was used to calculate the analytical inverse Fourier transform (IFT) which determines the Fourier transform limit (FTL) (the shortest pulse that could be supported by the bandwidth) of the laser spectrum. Signals that fell below 1% of the peak signal in the measured spectrum were set to zero and padded with zeros. The full width at half maximum (FWHM) of the resulting IFT was rounded to the nearest femtosecond, and determined to be ∼ 31 fs. Therefore, this is the shortest possible pulse that could be supported by our laser spectrum, representing a pulse with no spectral phase distortion. A Gaussian spectrum (80 nm FWHM) representing a compressed laser pulse with a Gaussian shape and FWHM of 31 fs was used as an analytical model for exploring the effects of GDD and TOD in all analytical simulations [e.g. Figs. 2(d)-2(e), 3 and Supplementary Fig. S2].

The effects of GDD on model laser pulses with and without the measured TOD was determined by using the open source python package PyNLO, which has the capability of analytically modeling the laser pulses with arbitrary spectral phase^15^. The effects of TOD on pulse compression was determined by adding the measured TOD from our laser source to the analytical Gaussian model pulses (∼ 31 fs). Then, to emulate pulse compression, these pulses (with fixed TOD) were given differing amounts of GDD in the range −1000 fs^2^ to +1000 fs^2^ [e.g., Fig. 2 (d)]. The effect of TOD alone on the ideal pulse profile (which implies all GDD has been removed) was explored by adding exactly the measured amount of TOD to the model pulse, and compared to the model pulse itself [e.g., Fig. 2 (e)]. Similarly, the effects of scaling TOD on pulse duration were demonstrated by adding a range of TOD values from 0 to 50,000 fs^3^ to our model pulses and recording their FWHM pulse duration. Simulated autocorrelation measurements were performed by analytically autocorrelating the temporal intensity envelope of the model pulses and subsequently deconvolving the autocorrelation by a Gaussian deconvolution factor 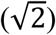. This simulates an intensity autocorrelation measurement in which the underlying pulse shape is unknown, but assumed to be Gaussian. To explore the effects of TOD on energy distribution in a pulse, we also analytically simulated the temporal intensity envelope with a range of TOD values [e.g., Fig. 3].

**Fig. 3.**
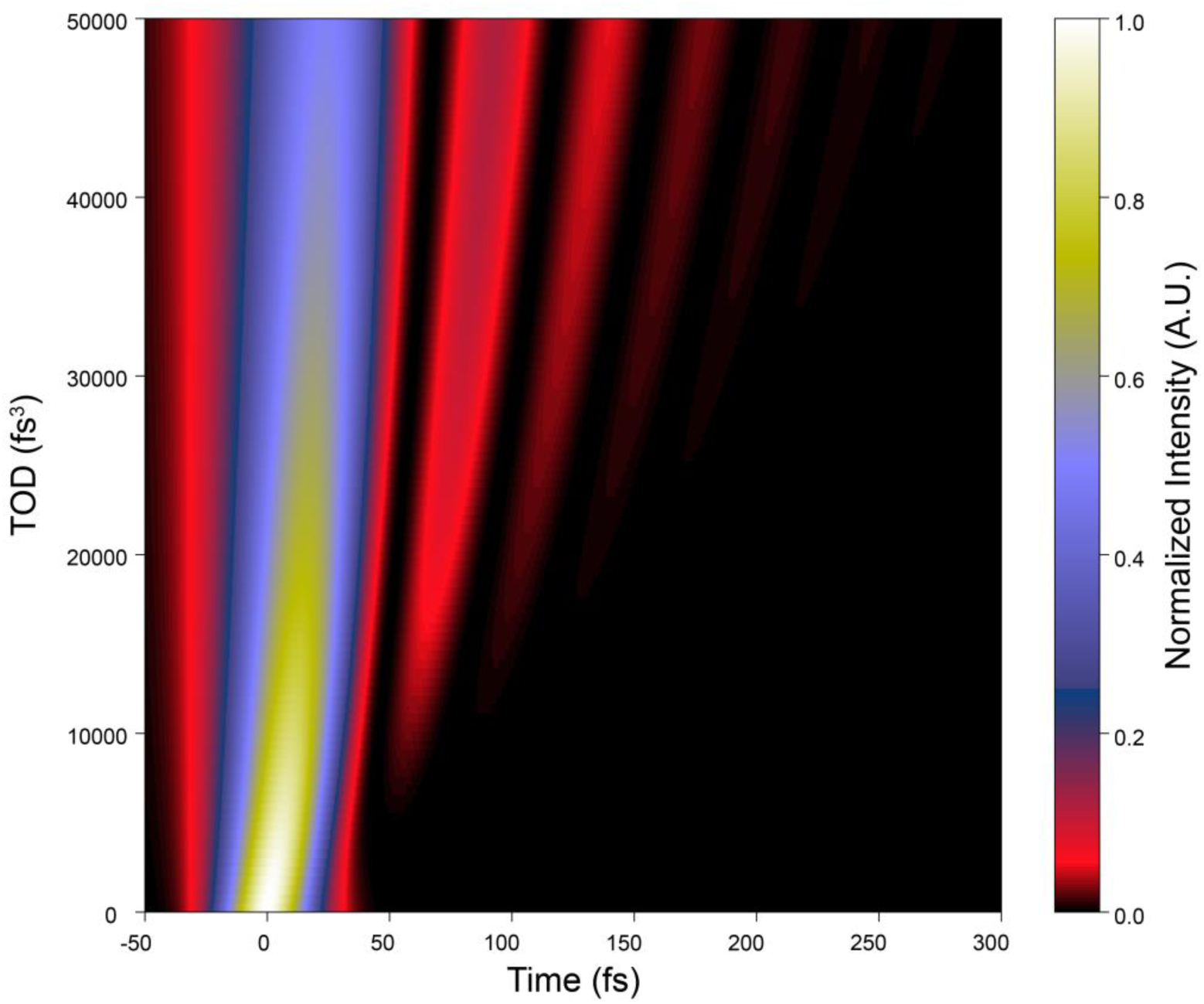
Effects of TOD on temporal energy distribution in a GDD compensated laser pulse. For our model pulse which was 31 fs FTL, the temporal intensity envelope on the horizontal axis is shown for a range of TOD values on along the vertical axis, assuming no other phase distortions, i.e., GDD = 0 fs^2^. Here, pulse energy is held constant as TOD is increased to visualize its effect on peak intensity at a given pulse energy.

### 2.6 Microscope imaging

Imaging was performed with a commercial multiphoton microscope (Investigator, Bruker Nano), which we further customized for three-photon imaging. The idler output (tuned to 1300 nm central wavelength) from a non-collinear optical parametric amplifier (NOPA-VISIR, Spectra Physics) pumped by a 1 MHz 70 W laser (Spirit 1030-70, Spectra Physics) was used as the three-photon excitation source. This two-stage optical parametric amplifier is the same design as that used in other commercial laser systems for three-photon imaging such as the Coherent Monaco-OPERA pumped by a different source laser. All imaging of fluorescent slides was performed using a 25× objective lens (XLPLN25XWMP2, Olympus). The dispersion of laser pulses was compensated by a prism compressor in the optical path before the microscope. The emitted photons were transmitted through a short-pass primary dichroic mirror (t770spxrxt, Chroma) and further filtered by a short-pass filter (ET770sp-2p-1500IR, Chroma) before being reflected by long-pass dichroic mirror (T565lpxr, Chroma) and ultimately transmitted through a band-pass filter (525 ± ?25 nm band-pass, Chroma) before being collected by a photomultiplier tube detector (H10770PB-40 SEL, Hamamatsu).

### 2.7 Deep-memory diode imaging

Laser pulses from the 1 MHz Spirit-NOPA tuned to 1300 nm central wavelength were attenuated by a half-wave plate (05RP02-40, Newport) and SF2 polarizing beam splitter (05FC16PB.9, Newport) to decrease the pulse energy, as needed. Laser pulses were then focused with a 16X microscope objective (N16XLWD-PF, Nikon) onto the surface of a GaP photodiode (DET25K2 or PDA25K2, Thorlabs) to induce three-photon absorption signals [Supplementary Fig. 3(a)]. For one-photon DMDI, the measurement does not require a focusing objective lens, and in this case an InGaAs photodiode was used (DET05D2, Thorlabs) [see Supplementary Fig. 3(c)].

Laser power was adjusted to achieve similar three-photon signal strengths in each category of the DMDI measurements of stable pulses, microsecond-scale fluctuations and millisecond-scale variations [see Fig. 5]. An example raw voltage trace from the GaP photodiode trace is included and was achieved at an incident average power of 18.4 mW [Supplementary Fig. S4].

**Fig. 4.**
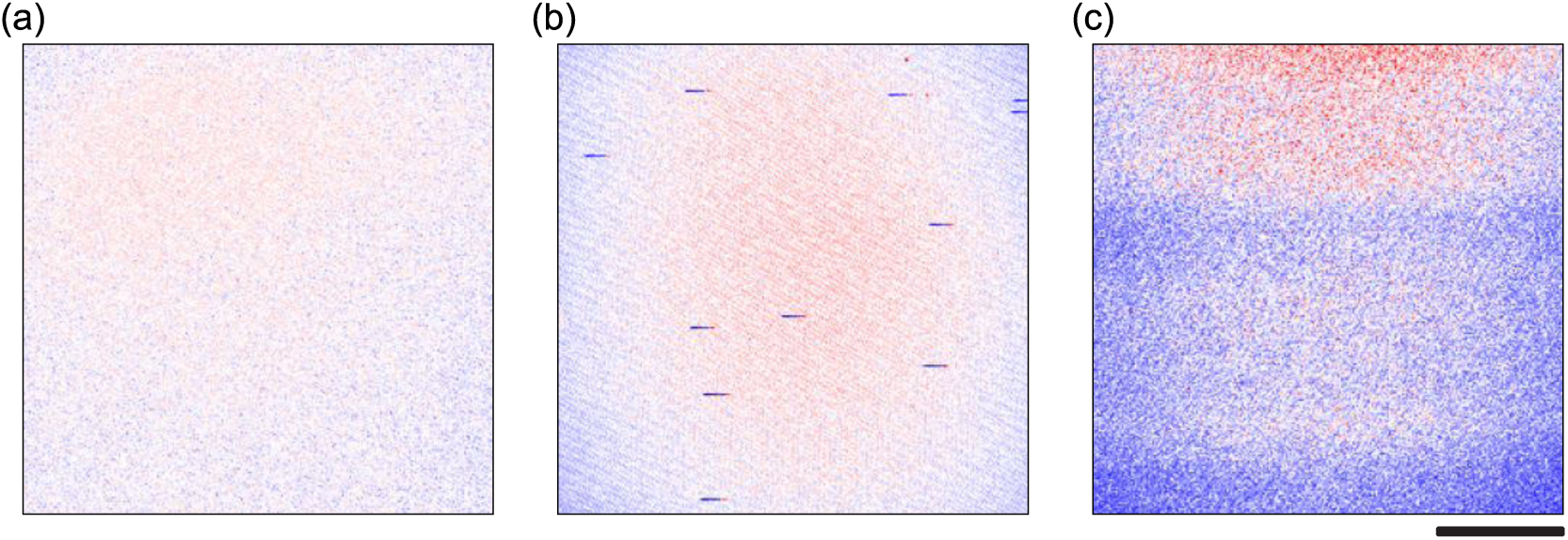
Three-photon microscopy images of a uniform fluorescent slide under normal conditions and in the presence of brightness artifacts. (a) Uniformly illuminated microscope image when the laser excitation source was optimized. In this and subsequent panels, minimum and maximum values of fluorescence in the image is scaled to ± 95% of the mean such that red represents signals that are higher than the mean, white are at the mean, and blue are below the mean. (b) Image with intermittent microsecond-scale ‘streaks’ of reduced brightness (dark blue pixels) followed by a very short rebound (red pixels). (c) Image with slower millisecond-scale temporal modulation of brightness. All images are 256 × 256 pixels, where each pixel represents the combined fluorescence signal from three laser pulses, i.e., an integration time or pixel dwell time of 3 μs from a laser with a repetition rate of 1 MHz. Scale bar 100 μm.

**Fig. 5.**
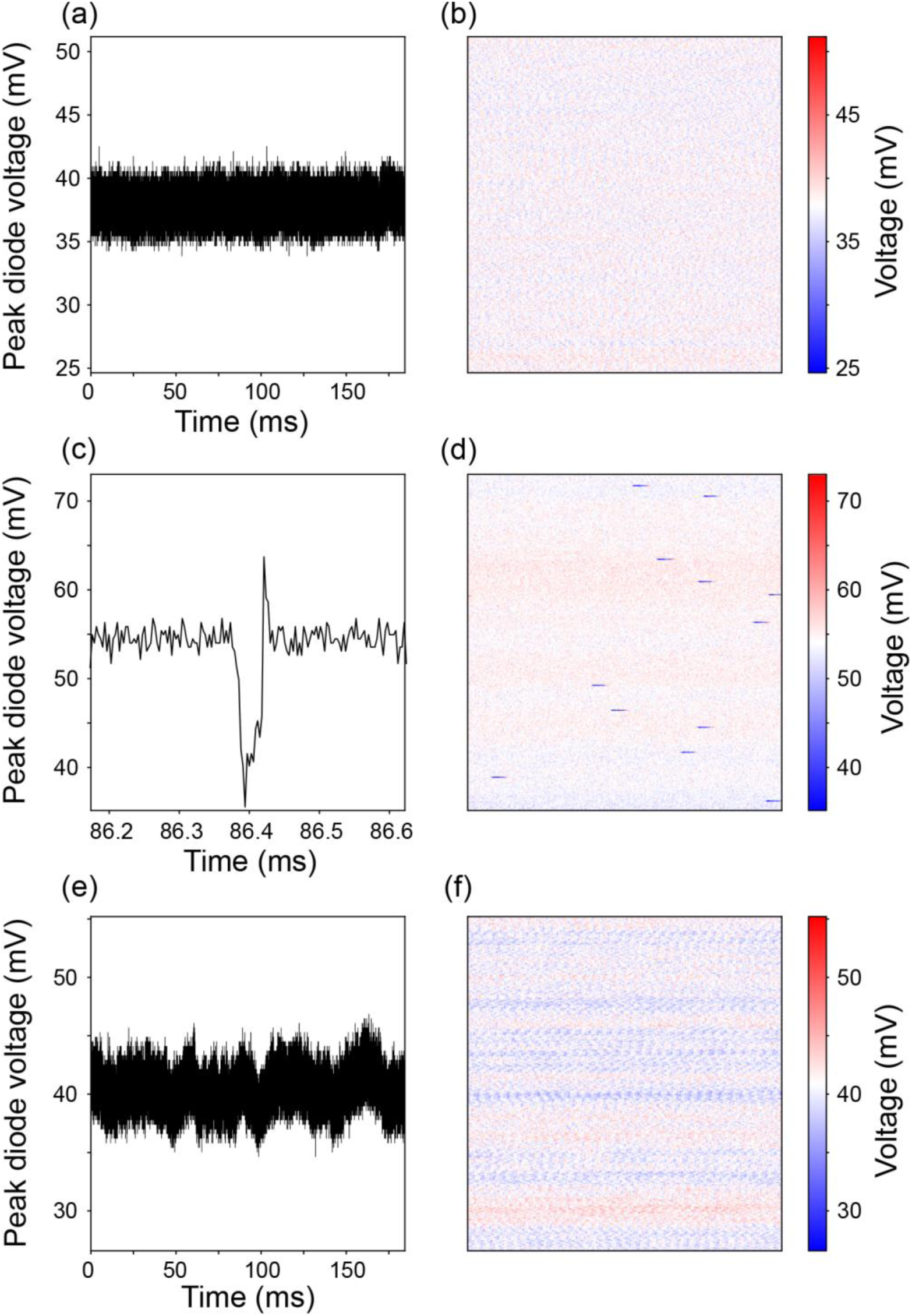
Direct measures of laser pulse intensity fluctuations using a photodiode. (a,b) Normal condition with stable laser pulses. (c,d) Microsecond-scale fluctuations of laser pulses with clear ‘streaks’ of reduced brightness. (e,f) Millisecond scale ‘wave’-like fluctuations. For each of the three conditions, to mimic a pixel integration time of 3 μs that was used during imaging (as shown in Fig. 4), here the photodiode data are arranged as peak voltage signals which were summed from every three successive pulses (left column). Then the same voltage signals are shown as 256 × 256 pixel images to visualize laser pulse intensity fluctuations in a manner akin to a microscope image (right column). The time axis in panels c and e are adjusted to reveal the relevant fluctuation patterns. The y-axis and color bar in each case are scaled to 35% of the mean voltage.

The photodiode signals were recorded by a high buffer memory oscilloscope (Picoscope 6404 or Picoscope 3406 DMSO, Picotech) at 62.5 megasamples per second or higher. When using the GaP photodiode and a 50 Ω terminating resistor, the rise and fall time of the voltage transient was ∼ 55 ns. This allowed for 7 or more samples per diode transient. The peaks in the diode voltage traces were identified and every three signal peaks were summed and plotted two ways [see Fig. 5]. Summing three successive voltage peaks from the photodiode measurements was only done so as to mimic the 3 μs integration time we use in microscope imaging experiments.

### 2.8 RMS% laser stability

The root-mean-square (RMS) is defined as Eq. 2,

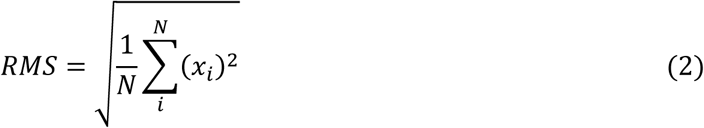

where *N* is the number of measurements and *x*_*i*_ is an individual measurement. On the other hand, the RMS% is defined as Eq. 3,

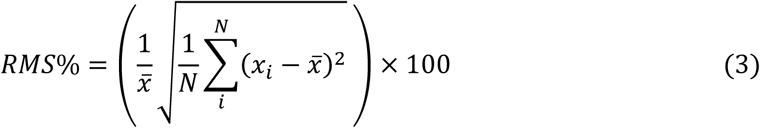

where 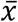 is the mean of all the measurements. RMS% is similar to the coefficient of variation and is readily identified as the ratio of the standard deviation to the mean, expressed as a percentage.

## 3 Results and Discussion

### 3.1 Factors that limit obtaining the shortest possible laser pulse

Due to their large bandwidth, laser sources used for three-photon microscopy are more challenging to harness than those used for two-photon microscopy. The bandwidth of laser pulses determines the shortest ideal pulse that can be achieved, known as the FTL, where larger bandwidths provide the potential for shorter pulses. The FTL is achieved when the peaks of the constituent wavelengths that make up the bandwidth of the laser pulse are coincident. If these constituent wavelengths have a nonzero phase shift with respect to each other, the laser pulse they construct will be distorted from its ideal shape. The phase of the constituent wavelengths (spectral phase) can be mathematically approximated by a polynomial, where the first two terms that affect the redistribution of pulse energy are group delay dispersion (GDD) and third order dispersion (TOD) (see Methods, Eq. 1). Laser pulses with larger bandwidths are more sensitive to phase distortions than laser pulses with smaller bandwidths. For example, an ideal 100 fs Gaussian laser pulse (typical for two-photon excitation) with ∼ 1000 fs^2^ of GDD would broaden to ∼ 104 fs. Conversely, an ideal 30 fs Gaussian laser pulse (typical for three-photon excitation) with ∼ 1000 fs^2^ of GDD would more than triple to ∼ 97 fs. Because the three-photon absorption process is more sensitive to changes in peak intensity, the larger susceptibility to broadening of the three-photon pulse will have a more dramatic effect on imaging when compared to laser pulses intended for two-photon excitation. Therefore, in three-photon applications, more attention must be paid to these phase distortions that could dramatically change the pulse properties.

### 3.2 A more accurate measurement of laser pulse properties for three-photon microscopy

#### 3.2.1 Simulated effects of phase distortions on laser pulses

Based on the above-mentioned theoretical principles that govern pulse duration, we first simulated the pulse distortion effects of GDD and TOD. Laser pulses at the FTL display an ideal pulse shape, where the instantaneous wavelength is fixed and the temporal intensity envelope is minimized [Fig. 1(a), 1(e)]. Pulses with GDD show a change in the instantaneous wavelength as a function of time and a broadened temporal intensity envelope [Fig. 1(b), 1(f)]. This phase distortion can be compensated with pulse compressors consisting of prism or grating pairs to approach the FTL. However, pulses with TOD show an asymmetric beating of the electric field and modulated temporal intensity envelope [Fig. 1(c), 1(g) and Fig. 1(d), 1(h)]. Such asymmetric distortions are more challenging to correct. Furthermore, TOD can be difficult to detect and is impossible to quantify when using intensity autocorrelations since they cannot uniquely determine the laser electric field or the intensity profile of laser pulses^16^. For example, the simulated intensity autocorrelations of Gaussian laser pulses with TOD could be incorrectly interpreted as pulses with no underlying phase distortions but a slightly different underlying pulse shape [Fig. 1(i)-1(l)]. Techniques like SHG-FROG are designed to quantify phase distortions missed by autocorrelation (see Methods). Specifically, the SHG spectrogram can be used to reconstruct the electric field and intensity profile of the underlying laser pulse [Fig. 1(m)-1(p)]. Ideal laser pulses at the FTL and laser pulses with GDD have SHG spectrograms with an elliptical shape, where an increase in temporal width is indicative of pulse broadening due to GDD [Fig. 1(m), 1(n)]. Most critically, in the presence of TOD, the SHG spectrogram has an hourglass shape, where adding GDD leads to an asymmetric broadening of the hourglass shape [Fig. 1(o), 1(p)].

#### 3.2.2 A comprehensive measurement of laser pulse properties

To directly measure the phase in our laser pulses, we built a SHG-FROG^13^ [see Methods and Supplementary Fig. S1]. The SHG-FROG measurements in Fig. 2(a) and Fig. 2(b) show a spectrogram and temporal pulse shape qualitatively consistent with the presence of GDD and TOD as observed in the simulations shown in Fig. 1(h) and Fig. 1(p), respectively. To quantify the amount of GDD and TOD, the measured spectral phase can be fit with a polynomial expanded about the central frequency [dashed black line in Fig. 2(c and see Methods, Eq. 1]. The second order term is used to calculate the GDD, which for our laser system was ∼ −590 fs^2^. The third order term is used to calculate the TOD, which for our laser was ∼ 22,750 fs^3^. Because the goal was to estimate the amount of TOD in the laser source, we performed these measurements at the laser head [see Supplementary Fig. S1] rather than after passing through the table optics, microscope, and objective lens, which would add more TOD^17^.

### 3.3 The consequences of laser TOD for three-photon imaging

#### 3.3.1 The effect of laser TOD on obtaining the shortest laser pulses

We added and removed GDD from a model Gaussian laser pulse containing TOD in the amounts measured from our laser source and compared these profiles to a model Gaussian pulse with no TOD [Fig. 2(d)]. When all GDD is removed from the laser pulse with TOD, it can only be compressed to ∼ 43 fs, as opposed to ∼ 31 fs if there was no TOD. Furthermore, in the presence of TOD, the duration of this pulse from autocorrelations deviate strongly from the actual pulse duration [compare open and filled red data points in Fig. 2(d)]. Whereas, in the absence of TOD, the pulse durations would otherwise match identically [see blue data points in Fig. 2(d)]. Thus, the ability to compress laser pulses by conventional methods and the accuracy of the intensity autocorrelation in determining pulse duration both deteriorate as a function of TOD. In essence, this deterioration is due to the asymmetric redistribution of energy in the laser pulse [see Supplementary Fig. S2].

#### 3.3.2 The effects of laser TOD on fluorophore brightness and heating of the sample

The leftover asymmetric redistribution of energy due to TOD (described above) will limit the achievable peak intensity and therefore imaging brightness in three-photon imaging. Laser pulses with our measured TOD versus no TOD (both with the same pulse energy) were analytically modeled and the case with TOD shows a decrease in peak intensity by ∼ 32% when compared to the case with no TOD [Fig. 2(e)]. In the instance of actual three-photon imaging, where laser pulses traverse through the rest of the path (table optics and microscope) the pulse would accumulate even more TOD, which would further limit their peak intensity even after all GDD was removed [see Fig. 3]. This implies that laser pulses with large TOD will need to operate at a higher average power to achieve the same brightness, when compared to laser pulses without TOD. Higher average power, even if available, comes with additional detriments of potential heat damage which will limit the duration of imaging and/or the penetration depth. Three-photon microscopy users attempting to maximize fluorescence brightness and minimize collateral heating^18-20^ will benefit most from systems that have laser pulses with minimal third (and higher) order phase distortions. The detrimental effects of these distortions can be minimized by utilizing laser sources without an excessive bandwidth (i.e., only enough bandwidth to reach the desired FTL) and can be managed by using optics designed to compensate specific phase profiles (i.e., with chirped mirrors).

Assuming phase distortions such as TOD are minimized such that the most efficient fluorescence excitation can be achieved, end users employing three-photon imaging in their laboratories must still contend with pulse-to-pulse intensity fluctuations that may exist in the excitation laser pulses. Therefore, in the following section we propose a new technique for evaluating the energy and intensity stability of laser pulses over timescales relevant for three-photon imaging.

### 3.4 Diagnosing pulse-to-pulse intensity fluctuations and visualizing their effects in imaging

#### 3.4.1 Imaging at the microscope

Several types of fluorescence artifacts on timescales relevant for imaging may be present in laser systems used for three-photon microscopy. Ideally, three-photon microscopy images of a uniformly fluorescent slide should present a field of view with relatively uniform brightness [Fig. 4(a)]. However, when artifacts on the order of ∼ 10–100 μs are present, they can manifest as short dark streaks in the microscope images [Fig. 4(b)]. Alternatively, when artifacts on the order of ∼ 10–100 ms are present in the microscope images, they can manifest as horizontal bright and dark banding [Fig. 4(c)]. The laser source used to generate each of the images in Fig. 4 was within specification with respect to RMS% average power stability (RMS% < 2%). However, RMS% power stability measurements use thermopile sensors, which typically have a time resolution of ∼ 1 s. Thus, these measurements give no information on variations in pulse-to-pulse energy or intensity over the time scale of microseconds to milliseconds, which are the critical timescales for three-photon imaging.

#### 3.4.2 A quantitative strategy to monitor laser pulse-to-pulse stability

To test the pulse-to-pulse intensity stability of the laser, we developed a technique called deep-memory diode imaging (DMDI) (see Methods). In DMDI, pulsed laser light is projected onto a photodiode and the photodiode signal is recorded by a high sampling rate oscilloscope with a large buffer to capture the pulse signal peaks at a high temporal resolution. To match the order of photo-absorption process in three-photon imaging, a tight-focusing element (e.g., a microscope objective) can be used to focus the laser onto a photodiode with an appropriately chosen band-gap that *disallows* one- and two-photon excitation [e.g., GaP; see Supplementary Fig. S3(a), 3(b)]. Like three-photon excited fluorescence, the photodiode signal recorded in this way is sensitive to peak pulse intensity and not just to total pulse energy. Alternatively, to simply capture the stability of pulse energy, a photodiode sensitive to one-photon excitation (e.g., InGaAs) may be used, which obviates the need for any focusing elements [Supplementary Fig. 3(c)]. To localize the source of any laser instability, the one- or three-photon photodiode being used in DMDI can be placed at different locations in the optical path, such as under the microscope objective [Supplementary Fig. S3(a)] or at the laser head [Supplementary Fig. S3(b), 3(c)]. Due to the use of high sampling rates and large buffer memory, DMDI can record relatively long and uninterrupted segments of pulse trains without aliasing of the signal [see Methods and Supplementary Fig. S4]. Therefore, DMDI offers a versatile and robust method to measure the laser pulse stability relevant for three-photon imaging.

#### 3.4.3 Linking photodiode measurements from the laser to microscopy images

We first used DMDI to monitor the pulse-to-pulse intensity stability of the laser by placing the three-photon photodiode (GaP) under the objective lens in our microscope [Supplementary Fig. S3 (a)]. Since the microscope images in Fig. 4 use a 3 μs integration time (and therefore each pixel represents the fluorescence signal of three laser pulses from a laser operating at 1 MHz repetition rate), the sum of three successive photodiode signal peaks was taken as a proxy for the brightness of one imaging pixel. This summed peak voltage signal was measured under conditions of artifact-free imaging as well as during imaging that had short and long timescale artifacts. Plotting the time courses of summed peak voltage signals under these three conditions revealed an underlying fluctuation in pulse intensities or lack thereof [Fig. 5(a), 5(c), 5(e)]. We then rearranged this linear array of summed peak voltages into a 256 × 256 matrix to resemble a three-photon image [Fig. 5(b), 5(d), 5(f)]. Whether the image is generated in the traditional manner with the aid of a microscope or by DMDI, the same phenomenological fluctuations are clearly identified. This is readily demonstrated in comparing Fig. 4(b) with Fig. 5(d), and Fig. 4(c) with Fig. 5(f), which were obtained when using laser sources that had 10–100 μs fluctuations, and 10–100 ms fluctuations, respectively.

#### 3.4.4 DMDI in laser diagnosis and repair

Beyond attributing the imaging artifacts to pulse intensity fluctuations, DMDI aided our laser manufacturer in improving pulse-to-pulse stability. To this end, we performed DMDI at the laser head using a one-photon photodiode, revealing that the 10–100 μs intensity fluctuations stemmed from fluctuations in pulse energy originating from the laser source itself (see Supplementary Fig. S5. The slower 10-100 ms intensity fluctuations may originate from several sources in addition to the laser, e.g., we found that air currents introduced in the beam path through small gaps (< 1 mm) in beam-shielding tubes can contribute to these fluctuations.

#### 3.4.5 A more sensitive statistic to describe laser pulse stability

Conventional statistics (e.g., RMS%, see Methods) on data collected either with thermopile sensors or photodiodes used in DMDI may fail to capture potential pulse-to-pulse instabilities. For example, the RMS% of the signal peaks of the pulse trains in all three conditions shown in Fig. 5 was 3–6%. We found that another simple statistic is more sensitive to large departures from the mean, i.e., 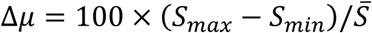, where *S*_*max*_ is the largest signal peak, *S*_*min*_ is the smallest signal peak, and 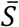 is the average signal peak over the entire time course of the measurement. When calculated on the photodiode pulse train data over three recording periods, this ‘min-max’ statistic (Δ*μ*) was 42.5±1.4% (for the stable condition) and 76.9±2.4% (for the 10–100 μs ‘streak’ artifacts). Statistics presented here are mean ± standard deviation. While RMS% measurements can capture large-scale oscillations in the amplitude variability of pulse trains, the ‘min-max’ statistic better captures the momentary 10–100 μs deviations from the mean. In addition to providing an intuitive snapshot of laser performance communicated through the ‘deep-memory diode image’, computing the ‘min-max’ statistic for data collected by DMDI provides a direct quantitative measurement of pulse-to-pulse stability over conventional methods like RMS%.

## 4 Future outlook

Although average power and pulse duration are measured to evaluate pulsed-laser performance as an industry standard, the equipment often used in these measurements (such as thermopile sensors and autocorrelators) do not capture the aspects of performance that are critical to three-photon microscopy. Due to few pulses being used for each imaging pixel and the cubic intensity dependence on fluorescence brightness per unit time, pulse-to-pulse intensity stability should be demonstrated in laser sources intended for three-photon microscopy (ideally using a technique like DMDI). Additionally, laser pulses should be shown to have minimal higher order phase using a spectral phase-resolved measurement (e.g., FROG), so that the laser pulses can be compressed and efficiently used in imaging. If pulse-to-pulse intensity stability is not demonstrated, three-photon imaging artifacts may be present. Finally, in laser sources that have large amounts of higher order spectral phase (e.g., TOD), laser pulses with more energy will be needed to achieve the same peak intensity, which will ultimately limit the depth and the duration of three-photon imaging in biological samples due to average power limits.

## Acknowledgements

We thank S.Aktürk for the source code we used for simulating SHG spectrograms and M Stanfield for computational support; the FROG group at Georgia Institute of Technology for their phase retrieval code; C.Xu for suggesting photodiode models; H.Jayakumar and A.Leikvoll for comments on the manuscript. This work was supported by grants from the NIH (R01 MH111447) and NSF (1707287).

## SUPPLEMENTARY INFORMATION

## Supplementary Figures

**Fig. S1.**
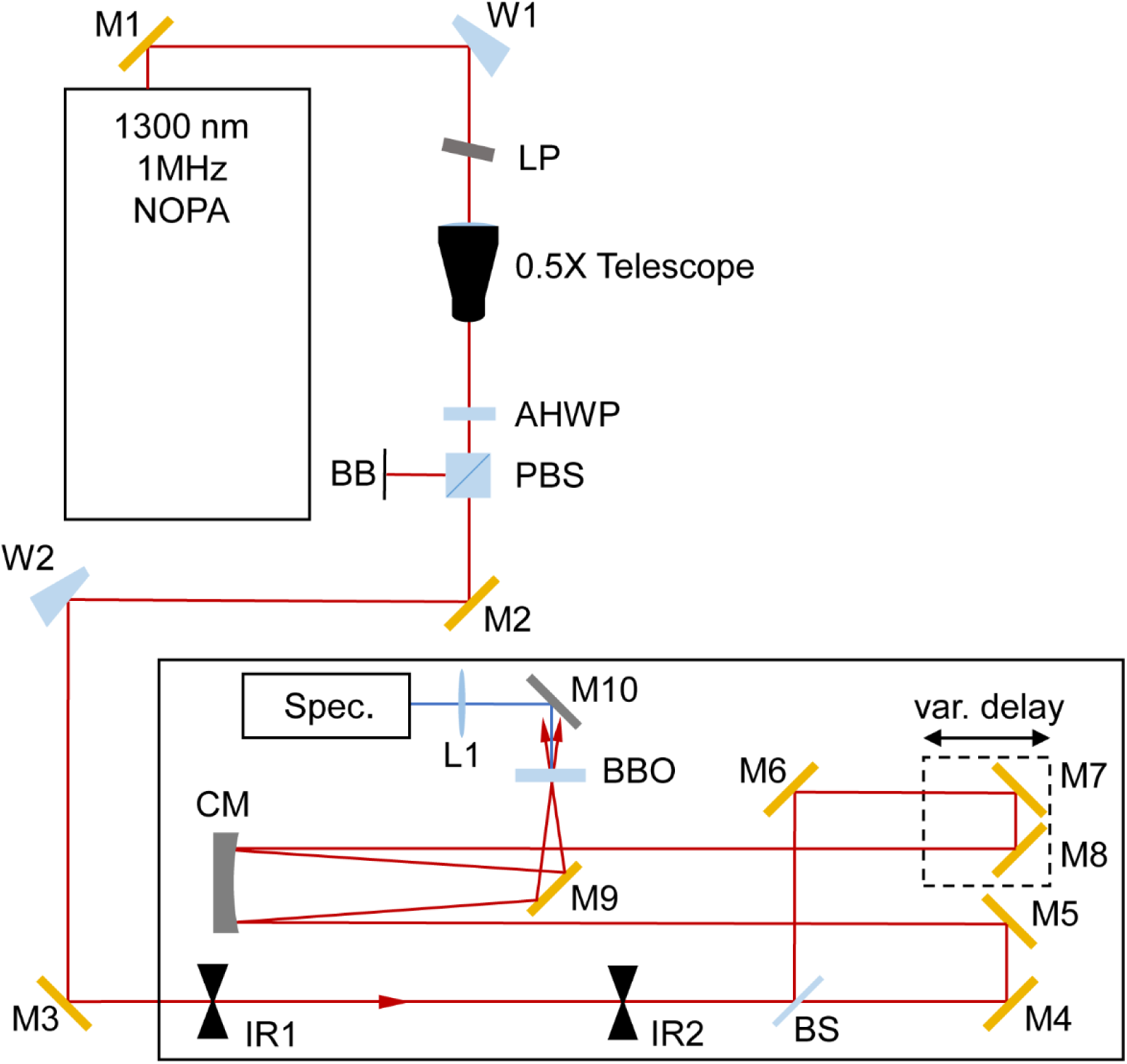
Schematic of light path and scanning SHG FROG setup. W1 and W2: uncoated fused silica wedges, LP: long-pass filter, AHWP: achromatic half-wave plate, PBS: polarizing beam splitter, BB: beam block, 0.5X Telescope: C-coated 0.5X down-collimating telescope, IR1 and IR2: irises, BS: pellicle beam splitter, CM: curved mirror, BBO: SHG crystal (β-Barium Borate), L1: BK7 lens, Spec.: Spectrometer, var. delay: retro-reflecting mirror pair mounted on a translation stage. Mirrors drawn as gold symbols are gold-coated and those shown in grey are silver-coated.

**Fig. S2.**
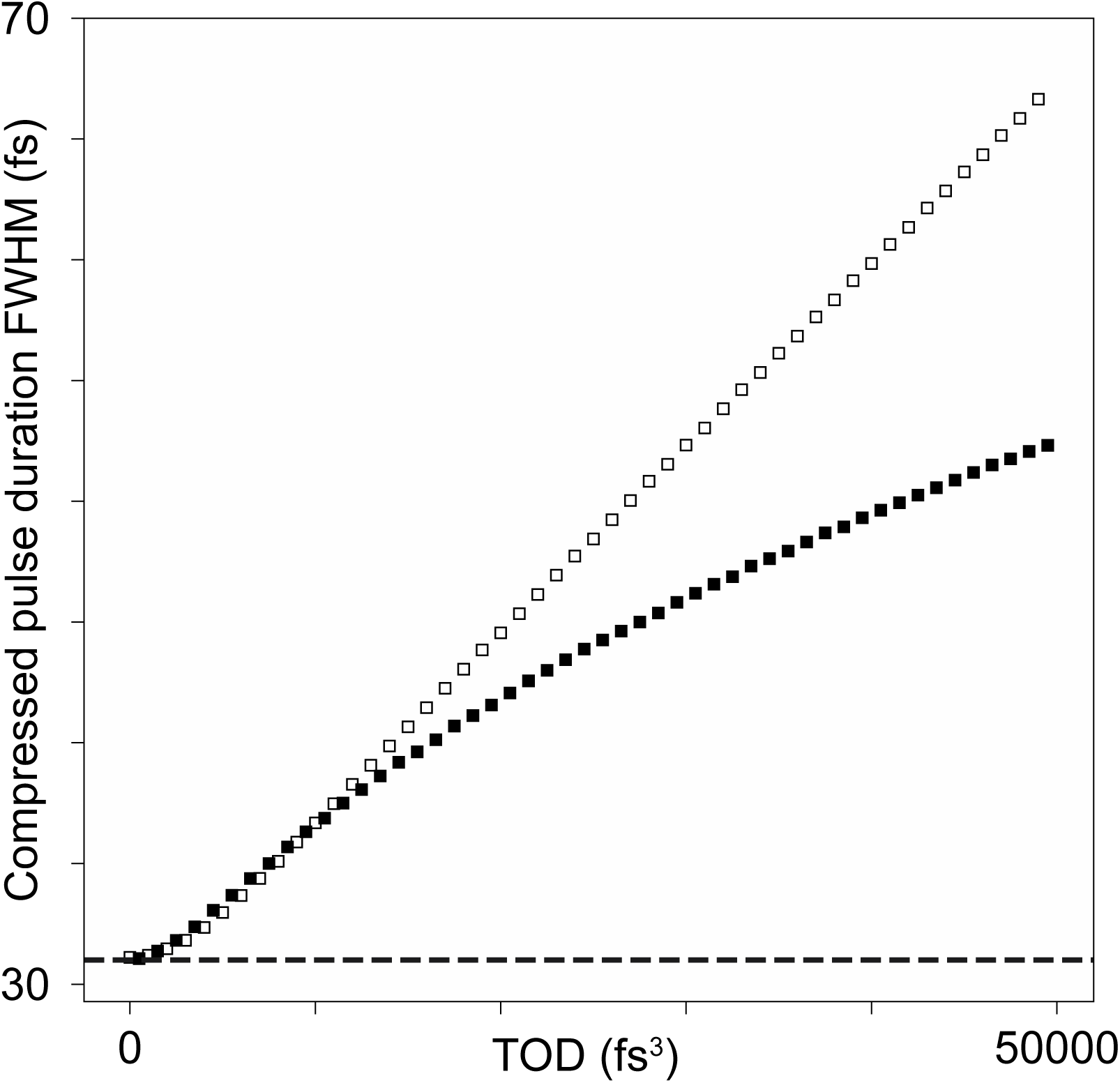
Limitations of using autocorrelations for measuring laser pulses that have TOD. Filled black squares represent the FWHM of the temporal intensity envelope of our model laser pulses (31 fs FTL) with a range of TOD values and no other phase distortions such as GDD. Unfilled black squares represent the pulse duration that would be measured for these same pulses by autocorrelations, if the underlying pulse shape was assumed to be Gaussian. The horizontal dashed line represents a perfectly compensated laser pulse with no phase distortions.

**Fig. S3.**
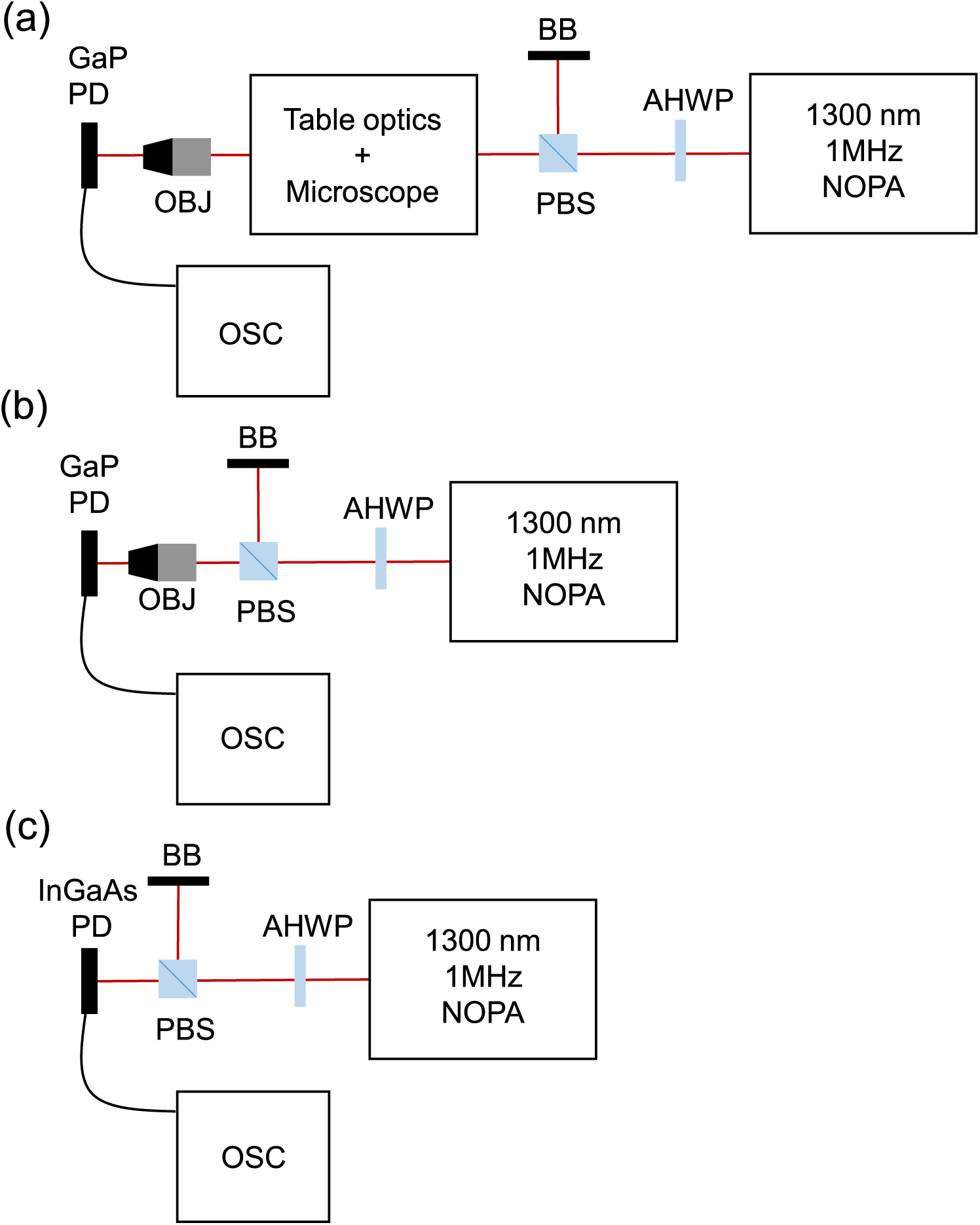
Setup for deep memory diode imaging (DMDI) from our laser source. (a) Three-photon photodiode recording using table and microscope optics. (b) Three-photon photodiode recording directly from the laser source with the exception of minimal components which were necessary to control laser power. (c) One-photon photodiode recording setup from the laser source, which by definition, does not require an objective lens to focus the light on the sensor. PD: photodiode, BB: beam block, PBS: polarizing beam splitter, HWP: half wave plate, OSC: large memory buffer oscilloscope, NOPA: non-collinear optical parametric amplifier, OBJ: objective lens.

**Figure S4.**
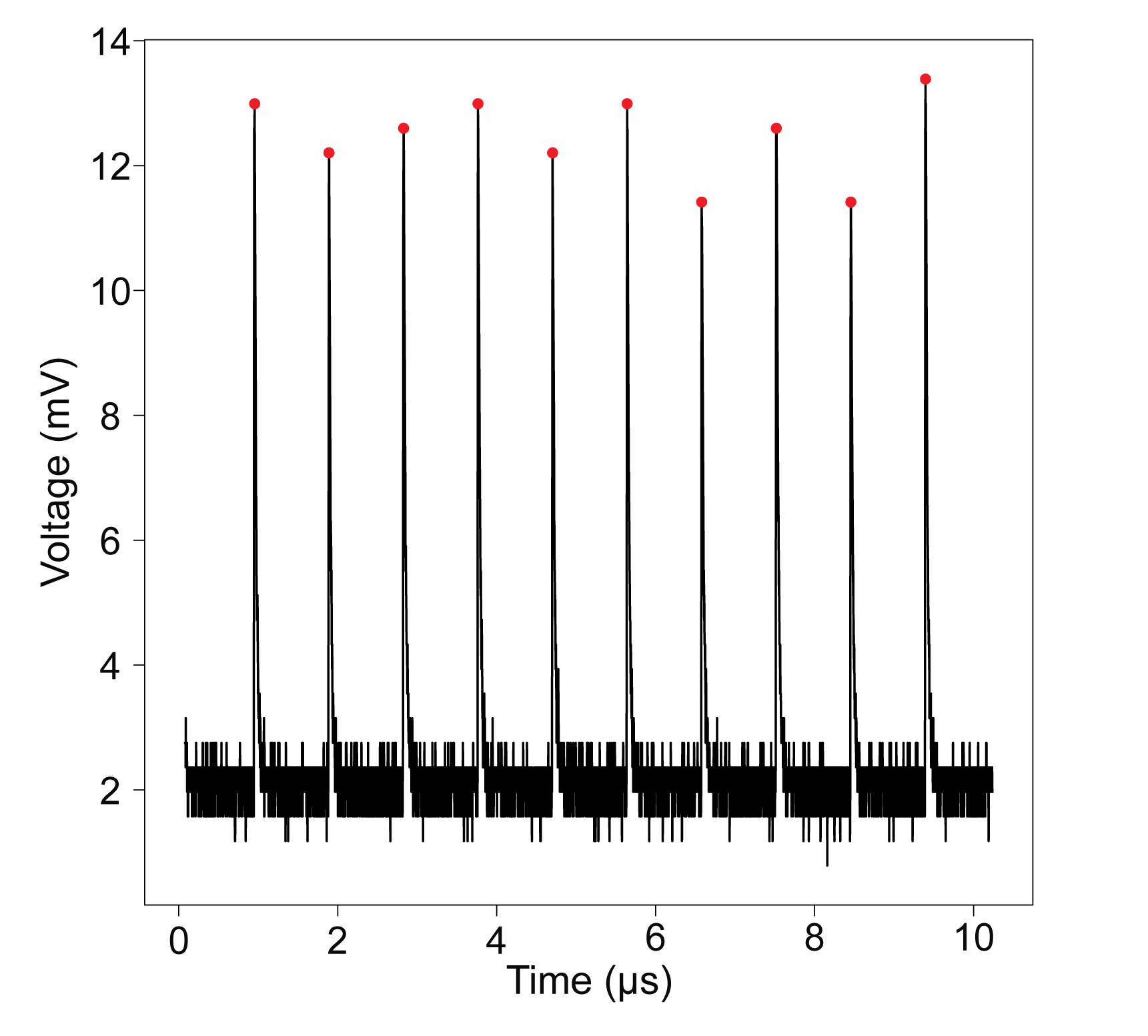
An example time course of a raw voltage trace of three-photon photodiode signals using our high bandwidth digital oscilloscope [see Methods and Supplementary Fig. S3 (a)]. Each red dot represents the peak voltage detected using the “find_peaks” function in the signal processing toolbox of the open source Python library SciPy.

**Fig. S5.**
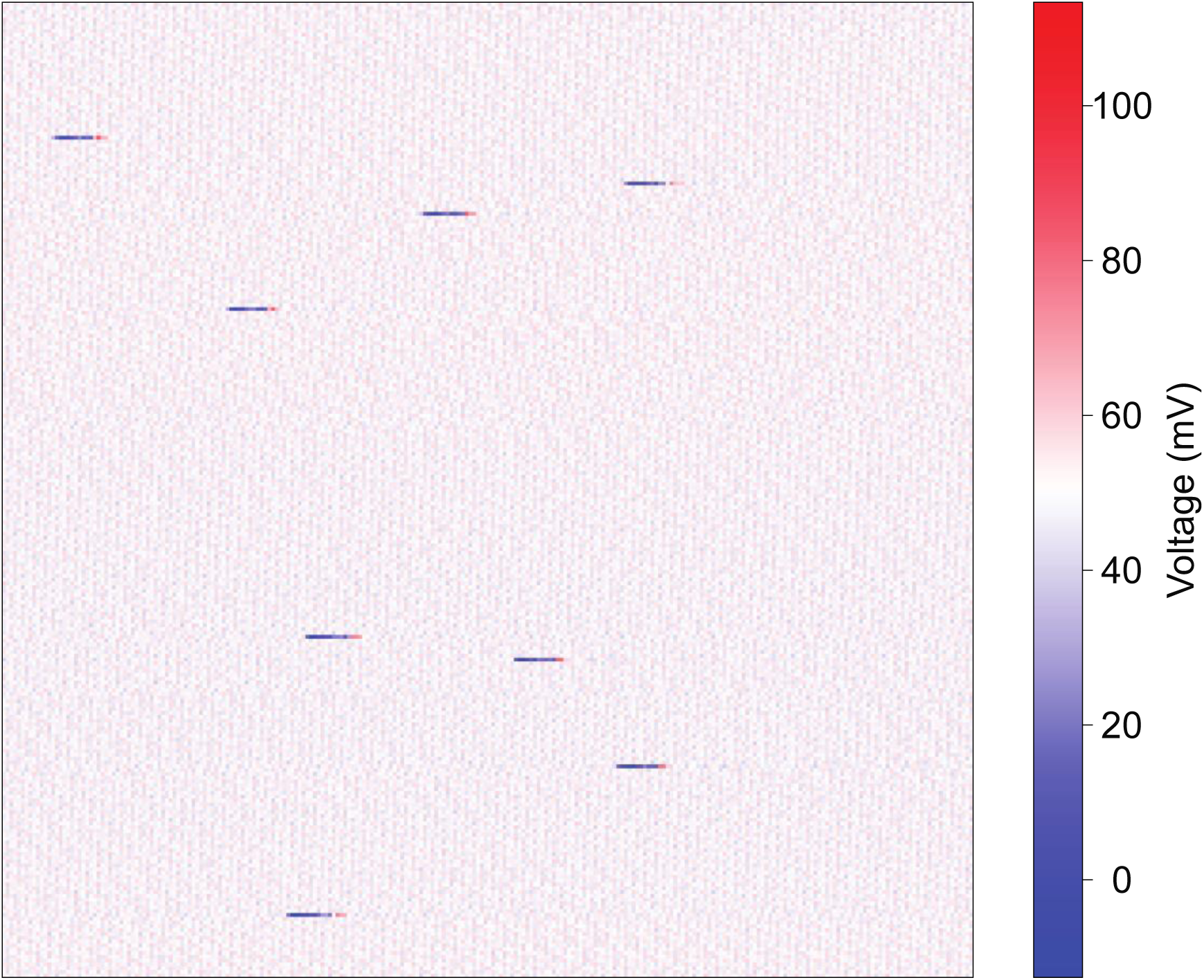
Sample DMDI measurement of laser instability using a one-photon absorption diode. Each pixel represents the sum of three voltage peaks. The 2D image representation is a 256 × 256 array of pixels to mimic a microscope image. Minimum and maximum values of the lookup table were scaled to ±125% of the mean such that red represents voltage peaks that are higher than the mean, white represents peaks that are at the mean, and blue represents peaks that are below the mean.

## List of Abbreviations

GDD: Group delay dispersion
TOD: Third order dispersion
DMDI: Deep-memory diode imaging
FTL: Fourier transform limit
SHG FROG: Second harmonic generation frequency resolved optical gating
FWHM: Full width at half maximum

